# Cerebral Microvascular Density, Permeability of the Blood-Brain Barrier, and Neuroinflammatory Responses Indicate Early Aging Characteristics in a Marfan Syndrome Mouse Model

**DOI:** 10.1101/2024.06.30.601409

**Authors:** T Curry-Koski, L Curtin, M. Esfandiarei, Thomas T. Currier

**Affiliations:** College of Medicine-Phoenix, University of Arizona, Phoenix, AZ, USA; Barrow Neurological Institute at Phoenix Children’s Hospital, Phoenix, AZ, USA; College of Graduate Studies, Midwestern University, Glendale, AZ, USA; Faculty of Medicine, University of British Columbia, Vancouver, BC, Canada; Phoenix VA Health Care System, Phoenix, AZ, USA

**Keywords:** Marfan Syndrome, Microvascular Density, Blood-Brain Barrier, Neuroinflammation, Neuropathology, Premature aging

## Abstract

Marfan Syndrome (MFS) is a connective tissue disorder due to mutations in fibrillin-1 (*Fbn1*), where a *Fbn1* missense mutation (*Fbn1^C1039G/+^*) can result in systemic increases in the bioavailability and signaling of transforming growth factor-β (TGF-β). In a well-established mouse model of MFS (*Fbn1^C1041G/+^*), pre-mature aging of the aortic wall and the progression of aortic root aneurysm are observed by 6-months-of-age. TGF-β signaling has been implicated in cerebrovascular dysfunction, loss of blood-brain barrier (BBB) integrity, and age-related neuroinflammation. We have reported that pre-mature vascular aging in MFS mice could extend to cerebrovasculature, where peak blood flow velocity in the posterior cerebral artery (PCA) of 6-month-old (6M) MFS mice was reduced, similarly to 12-month-old (12M) control mice. Case studies of MFS patients have documented neurovascular manifestations, including intracranial aneurysms, stroke, arterial tortuosity, as well as headaches and migraines, with reported incidence of pain and chronic fatigue. Despite these significant clinical observations, investigation into cerebrovascular dysfunction and neuropathology in MFS remains limited. Using 6M-control (*C57BL/6*) and 6M-MFS (*Fbn1^C1041G/+^*) and healthy 12M-control male and female mice, we test the hypothesis that abnormal Fbn1 protein expression is associated with altered cerebral microvascular density, BBB permeability, and neuroinflammation in the PCA-perfused hippocampus, all indicative of a pre-mature aging brain phenotype.

Using Glut1 staining, 6M-MFS mice and 12M-CTRL similarly present decreased microvascular density in the dentate gyrus (DG), cornu ammonis 1 (CA1), and cornu ammonis 3 (CA3) regions of the hippocampus. 6M-MFS mice exhibit increased BBB permeability in the DG, CA1, and CA3 as evident by Immunoglobulin G (IgG) staining, which was more comparable to 12M-CTRL mice. 6M-MFS mice show a higher number of microglia in the hippocampus compared to age-matched control mice, a pattern resembling that of 12M-CTRL mice.

This study represents the first known investigation into neuropathology in a mouse model of MFS and indicates that the pathophysiology underlying MFS leads to a systemic pre-mature aging phenotype. This study is crucial for identifying and understanding MFS-associated neurovascular and neurological abnormalities, underscoring the need for research aimed at improving the quality of life and managing pre-mature aging symptoms in MFS and related connective tissue disorders.

## Introduction

Marfan Syndrome (MFS) is the most common monogenetic autosomal-dominant disorder of connective tissue due to mutations in the gene encoding for fibrillin-1 (*Fbn1*) that presents with manifestations in many organs and systems in the body, resembling phenotypes of premature aging.^1,2^ The *Fbn1* gene encodes for a large glycoprotein, a structural component of microfibrils in the extracellular matrix (ECM).^3^ Therefore, *Fbn1* mutations can result in a wide range of complications, particularly those associated with connective tissues that require microfibril scaffolding, such as elastin. Thus, the cardiovascular, musculoskeletal, and pulmonary systems are commonly affected, with ascending aortic aneurysm and rupture being the most life-threatening manifestation. MFS vascular complications are attributed to compromised ECM, medial elastin fiber fragmentation, and endothelial dysfunction, all leading to vascular wall stiffening and weakening, contributing to the vascular aging phenotype.^4–9^

Fbn1 protein acts by binding and sequestering the latent form of the cytokine transforming growth factor-β (TGF-β) via a latent binding protein (LTBP), forming a large sequestering domain. However, the abnormal and dysfunctional Fbn1 protein fails to sequester TGF-β in its inactive form. This uncontrolled release of bioavailable TGF-β facilitates the activation of downstream signaling molecules involved in inflammatory responses, including matrix metalloproteinases (MMPs), which contribute to the degradation of elastin, a key and crucial component of the ECM structure within the wall of elastic arteries.^7,8,10^ Indeed, previous studies in MFS patients and experimental mouse models have confirmed increased expression and activity of MMP-2/-9 in the peripherary.^5,7,8,11^

Cerebrovascular aging increases the risk for neurological deficits such as cognitive decline, dementia, stroke, and cerebral aneurysms.^12–14^ MFS patients and mice display accelerated and pre-mature aging phenotypes within the peripheral vasculature, including increased vascular wall stiffness, endothelial dysfunction, decreased smooth muscle contractility, mitochondrial dysfunction, increased inflammatory infiltrates, and elevated reactive oxygen species (ROS) production within the vessel wall.^7–9,15–17^ This pre-mature aging phenotype has been observed and supported in multiple systems of the body, including the integumentary and circulatory systems.^18,19^ MFS patients experience other age-related changes such as worsening joint pain and stiffness, vision problems, hearing loss, skin elasticity, and lung complications.^1,2,19–21^ MFS-associated symptomology are also implicated in pathophysiological manifestations related to aging, such as inflammation, ROS production, arterial remodeling, and elastolysis all seen in the periphery thus far. Publications from our lab further supported this pre-mature aging phenotype in cerebral vascular function in transgenic mice with a *Fbn1* mutation *(MFS mice)*. Specifically, we demonstrated that the peak blood flow velocity in the posterior cerebral artery (PCA) of 6-month-old male MFS mice is reduced, showing similarities to that of middle-aged, 12-month-old control male mice.^18^

Cerebral blood flow is a critical physiological process required to ensure the brain receives the necessary oxygen and nutrients to function properly. Microvascular density, the number of microvessels in a given area such as the brain, plays a key role in cerebral blood flow. For instance, microvascular density is integral to regulating cerebral blood flow by influencing capacity and efficiency of blood delivery to the brain.^22^ Factors influencing microvascular density include but are not limited to angiogenic factors, hypoxia, and vascular stress. Microvascular density in the brain is commonly assessed using histological methods, including staining of glucose transporter 1 (Glut1), a protein predominately expressed on endothelial cells of brain capillaries that facilitates glucose transport across the blood-brain barrier (BBB).^23,24^ Aging is associated with a reduction in microvascular density, contributing to decreased cerebral blood flow, which can lead to impaired neurological function.^22,25,26^ The MFS-associated premature aging phenotype and decreased cerebral blood flow velocity warrants further investigation to evaluate microvascular density in the brain.

The BBB is the first line of defense and a key player in maintaining homeostasis in the central nervous system. This highly selective and semipermeable barrier is made up of different cell types and proteins comprising the neurovascular unit. The neurovascular unit is composed of endothelial cells that line the lumen of the blood vessel and are interconnected by a series of tight junctions to ensure strict segregation of the cerebral and vascular environments. Breakdown of the tight junction components, claudins and occludins could lead to an increase in permeability. The basal lamina of the endothelial cells adds another degree of separation between the two environments. Pericytes also line the endothelial cells involved in regulating blood flow and maintaining the BBB.^27^ In addition, astrocytic end feet wrap around the vessel and further regulate the passage of different molecules, regulating the cerebral environment. Increased BBB permeability is a phenomenon associated with aging, leading to various theories about the pathogenesis of aging-related neuropathologies such as dementia, Parkinson’s disease, and mild cognitive impairment, suggesting a link between the compromised integrity of the BBB and the development of these neurological conditions.^28–30^ Vascular dysfunction has also been associated with increased risk for BBB permeability.^31,32^ In addition, increased MMPs -2/-9 expression has been linked to disruption of tight junctions and increased BBB permeability.^32–34^ It is important to note that MMP-2/-9 are found to be upregulated in the blood and aortic tissue of MFS patients and mouse models. This makes it plausible that their expression might also be elevated in the cerebrovasculature and brain of MFS mice, potentially contributing to the observed neuropathological changes.^34–37^

Microglia are the resident immune cells of the brain and are significantly involved in neuroinflammatory responses, phagocytosis, homeostasis, and regulation of synapses. Neuroinflammation is characterized by the activation of microglia, which is marked by changes in morphology and an increased number of Ionized Calcium-Binding Adapter Molecule 1 (iba-1)-positive cell bodies. However, excessive and chronic activation of microglia, as observed in aging and neurodegenerative diseases, can result in impaired microglial function and neuronal damage, leading to neurodegeneration and cognitive decline.^38,39^ Increased permeability of the BBB is a well-recognized factor contributing to neuroinflammation, creating a vicious cycle where neuroinflammation further increases BBB permeability.^40^ In addition, increased MMPs -2/-9 expression has been shown to exacerbate BBB permeability and stimulate neuroinflammation, further augmenting the cycle of damage and inflammation within the brain.^41^

While aortopathy in MFS patients and animal models has been extensively researched by numerous groups, our understanding of other manifestations of MFS remains significantly limited. Currently, there is a notable gap in preclinical investigation into these manifestations and their possible impact on neurological outcomes. This lack of research highlights the need for a broader exploration of MFS beyond its cardiovascular implications, to fully understand the syndrome’s comprehensive effects on health and quality of life. In the present study, we aimed to test the hypothesis that a missense mutation in the *Fbn1* gene (*Fbn1^C1041G/+^*) results in changes in neuropathology in the well establish MFS mouse model. Furthermore, we aimed to determine whether the outcomes observed in 6-month-old MFS (6M-MFS) mice more closely resemble the phenotypes seen in 12-month-old healthy control mice (12M-CTRL), highlighting the potential premature and early aging brain phenotype in the MFS mouse model.

## Methods

### Experimental Animals

Animal care was conducted according to the National Research Council Guide for the Care and Use of Laboratory Animals and the Guidelines for Animal Experiments and the Midwestern University Animal Care and Use Committee [IACUC protocols AZ-3006 & AZ-2936]. Mice were group-housed (up to 5 mice per cage) in a 12/12h light-dark cycle with food and water available *ad libitum*. Breeding pairs were obtained from Jackson Laboratory and backcrossed for 8 generations. A breeding colony of MFS mice was established and maintained at Midwestern University Animal Facility (IACUC breeding protocol AZ-2989). Mice used in this study were heterozygous for an *Fbn1* allele encoding a cysteine substitution in the epidermal growth factor-like domain in *Fbn1* to glycine (*Fbn1^C1041G/+^*), giving the mice the most common type of mutation seen in MFS patients presented with vascular dysfunction and aortic root aneurysm. This study utilized adult male and female *Fbn1^C1041G/+^*(MFS) mice at 6 months of age, as well as male and female *C57BL/6* (CTRL) mice at 6 and 12 months of age. MFS mice present established and evident vascular defects (e.g., aortic wall elastin fragmentation, aortic root aneurysm, and increased aortic wall stiffness) by 6 months of age. This time point is equivalent to 30-35 human years and was compared to middle-aged (12-month-old) CTRL mice, to evaluate the MFS-associated accelerated vascular aging phenotype. These MFS mice are bred on a *C57BL/6* background, making *C57BL/6* mice the appropriate controls for this study. After genotyping via tail snip and PCR testing, an N=4-5/group with combined male and female mice were randomly assigned to naïve 6M-old CTRL and MFS, and 12M-old CTRL mice for immunohistochemistry and utilized for all measures described. As the first known study to evaluate neuropathology in MFS mice, we included both male and female mice in order to evaluate these manifestations overall in MFS. Future investigations would benefit from including a larger sample size for both males and females in order to evaluate sex as a biological factor contributing to neuropathology. Pre-determined welfare exclusion criteria included removing any mouse from the study with visible wounds requiring veterinarian intervention or vocalization of pain that cannot be managed. No animals required exclusion due to welfare concerns.

### Immunohistochemistry

Mice were euthanized using 5% isoflurane inhalation followed by cervical dislocation, and brain tissue removed and hemisected (N=4-5/group). One hemisphere was post-fixed in 4% paraformaldehyde and sent to Neuroscience Associates (Knoxville, TN) for proprietary embedding techniques, sectioning, and staining of Glut1 to assess microvascular density, IgG to evaluate BBB permeability, and Iba-1 to visualize microglia for neuroinflammatory phenotypes. Brightfield 40x images of the hippocampus were analyzed using ImageJ to assess pixel density for Glut1 and IgG-stained sections. As previously described by our laboratory, the ImageJ skeleton Analysis plugin was utilized to assess microglial branch points, endpoints, branch length, and the number of microglia soma as a global indication of neuroinflammation for Iba-1-stained sections^42,43^ (**Figure 1**.). The hippocampus, perfused by the PCA, was targeted and analysis done in the DG, CA1, and CA3 of the hippocampus. Three images were taken per DG, CA1, and CA3, respectively, per coronal brain section, where three sections were assessed per animal, for a total of 9 images per animal averaged for each data point in each region of interest (27 images per animal).

**Figure 1.**
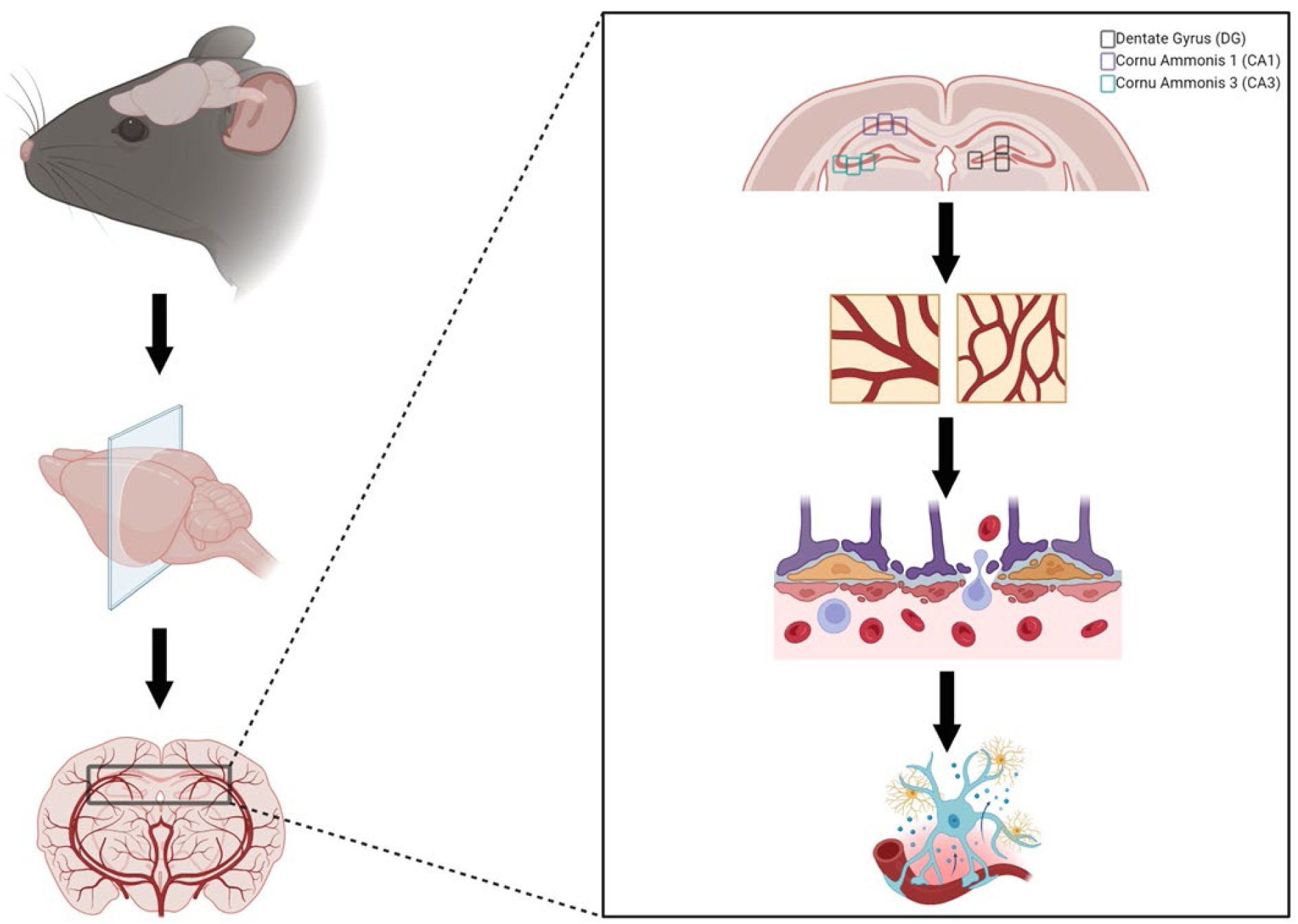
Representetion of neuropathotogy evaluation. Brains were collected, post-fixed in 4% paraformaldehyde, blocked, sectioned, and stained by Neuroscience Associates (Knoxville, TN). Neuropathology was evaluated by investigating Glut1 staining to evaluate microvascular density, lmmunoglobulin G (lgG) staining to assess blood-brain barrier permeability, and iba-1 staining to investigate neuroinflammation in three regions of interest within the hippocampus (dentate gyrus, cornu ammonis 1, and cornu ammonis 3).

### Statistical Analysis

All graphs and statistical significance were created and determined with GraphPad Prism software. The sample size for experimental groups was calculated based on desired endpoints where power was set at 90% and α=0.05 utilizing the Mean ± SD from our and other previously published data (N=4-5/group).^42^ One-tailed T-tests were performed on data sets comparing 6M-CTRL and 6M-MFS mice. One-way analysis of variance (ANOVA) test was performed on data sets with one independent variable (e.g., phenotype) comparing 6M-CTRL, 6M-MFS, and 12M-CTRL mice. Our assessment evaluated the phenotypic similarity and differences between these cohorts thus 6M-MFS and 12M-CTRL mice were evaluated for the independent variable of phenotype. Tukey post-hoc was performed on ANOVA tests. All data sets were tested for outliers utilizing ROUT (Q=1%) and, if determined, were excluded. Furthermore, data sets were evaluated for Gaussian distribution to test for normality of residuals using the Kolmogorov-Smirnov test. Sex differences were not evaluated due to low sample sizes. Thus, we could not make reasonable statistical interpretations on sex as a biological factor in this study. Significance was determined as a *P*-value ≤ 0.05. All *P*-values are reported in the graphs, and raw data can be accessed by contacting the corresponding author. Personnel were blinded to genotype, age, and sex during analysis. Secondary analysis was performed by blinded personnel to replicate and confirm findings.

## Results

### Hippocampal microvascular density is decreased by age and genotype

To investigate neurovascular alterations, we sought a method to evaluate gross morphological changes in hippocampal microvasculature. Glut1 staining was utilized to mark vascular endothelial cells, a well-known method for evaluating microvascular density (**Fig. 2A, 3A, 4A**). 6M-MFS mice demonstrated decreased microvascular density in the dentate gyrus (DG) of the hippocampus compared to age-matched control mice (**Fig. 2B**), yet no difference is seen when compared to 12M-CTRL mice (**Fig. 2C**). Microvascular density was decreased in 6M-MFS compared to 6M-CTRL (**Fig. 3B**), and 12M-CTRL mice compared to 6M-CTRL mice in the cornu ammonis 1 (CA1) (**Fig. 3C**). Microvascular density was decreased in 6M-MFS compared to 6M-CTRL (**Fig. 4B**), and 12M-CTRL mice compared to 6M-CTRL mice in the cornu ammonis 3 (CA3) (**Fig. 4C**), while no differences were seen between 6M-MFS and 12M-CTRL mice.

**Figure 2.**
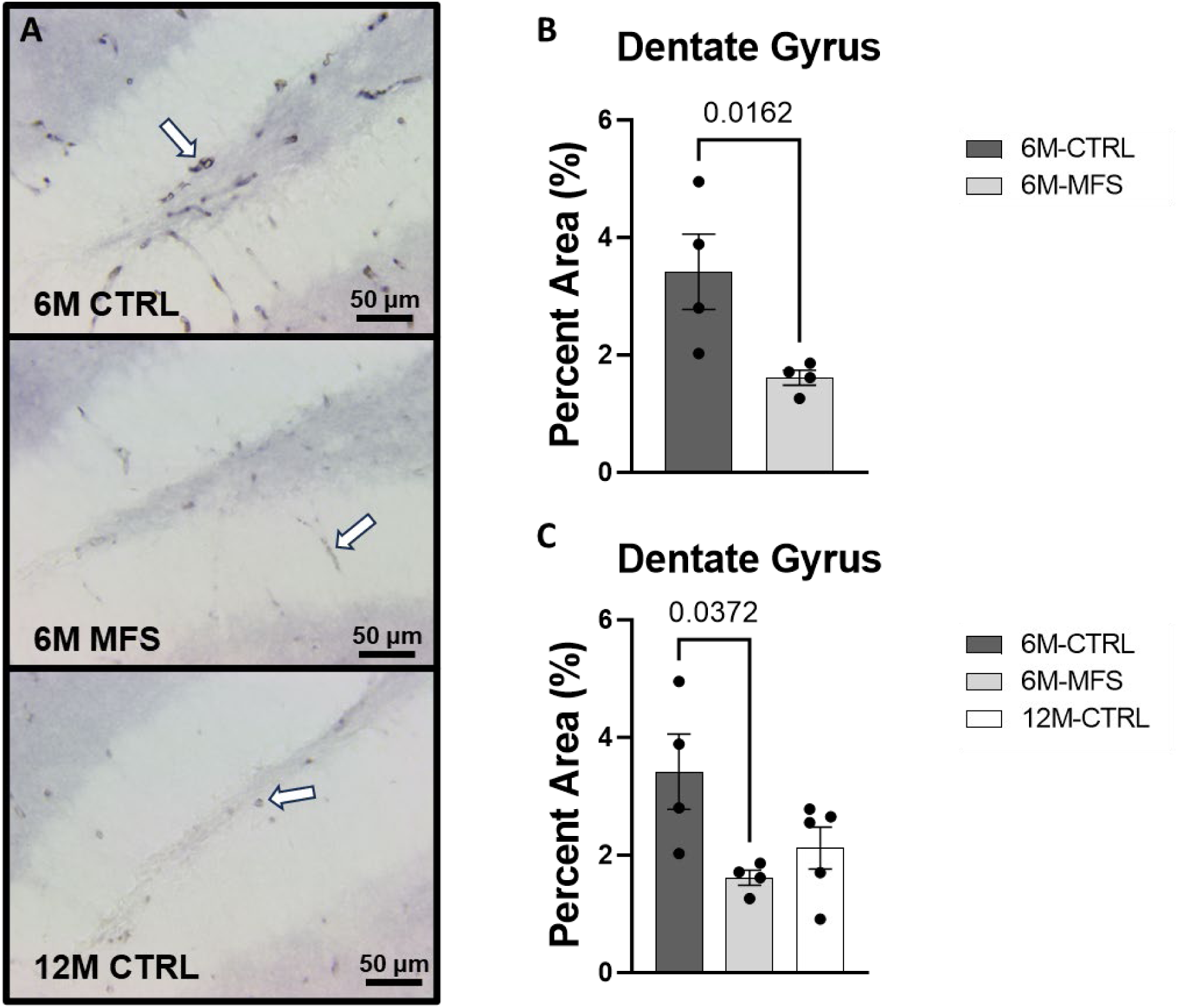
Glutl staining in the Dentate Gyrus (DG) is decreased as a function of genotype and age, indicative of decreased microvascular density. A) Representative images of Glutl staining in the DG of the hippocampus in 6M-CTRL, 6M-MFS, and 12M-CTRL respectively. Examples of positive signals are represented by white arrows. B) Glutl staining is decreased in 6M-MFS DG compared to 6M-CTRL, indicative of decreased microvascular density. C) 6M-MFS compared to 12M-CTRLmice demonstrate no significant differences in Glutl staining.

**Figure 3.**
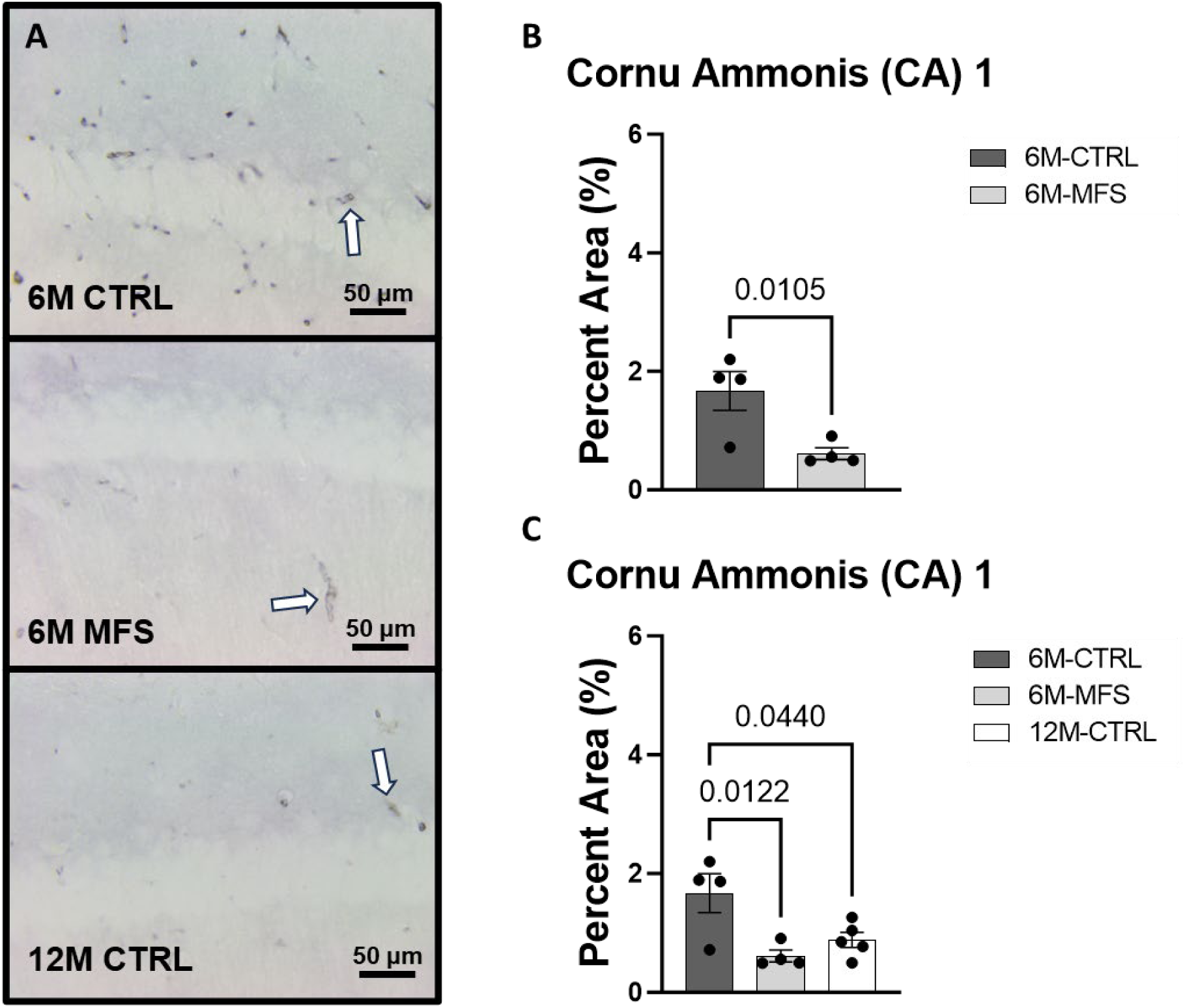
Glutl staining in the cornu ammonis 1 (CA1) is decreased as a function of genotype and age, indicative of decreased microvascular density. A) Representative images of Glutl staining in the CA1 of the hippocampus in 6M-CTRL, 6M-MFS, and 12M-CTRL respectively. Examples of positive signals are represented by white arrows. B) Glutl staining is decreased in 6M-MFS CA1 compared to 6M-CTRL. C) 12M-CTRLCA1 demonstrate decreased Glutl staining compared to 6M-CTRL, with no differences compared to 6M-MFS mice.

**Figure 4.**
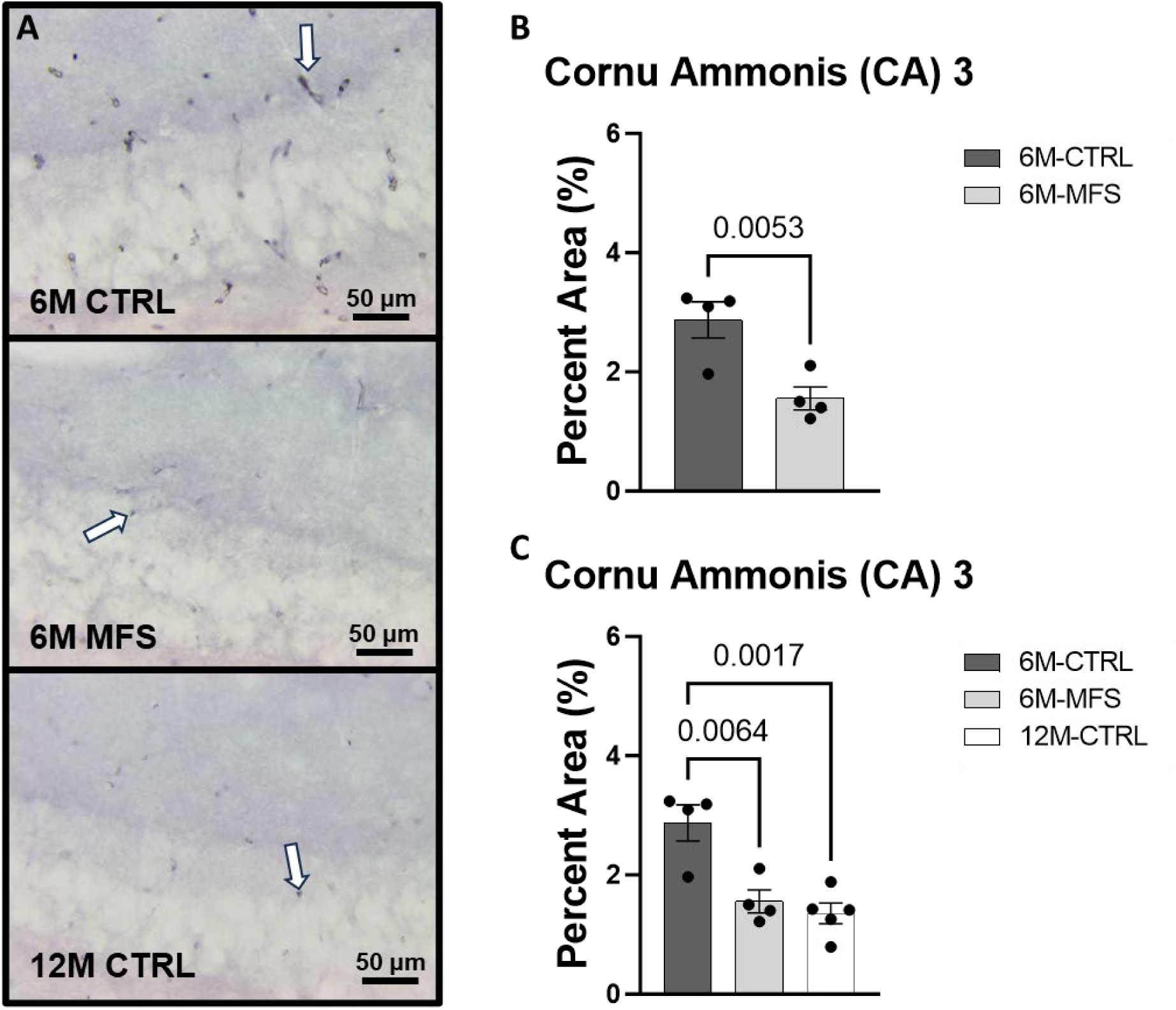
Glutl staining in the cornu ammonis 3 (CA3) is decreased as a function of genotype and age, indicative of decreased microvascular density. A) Representative images of Glutl staining in the CA3 of the hippocampus in 6M-CTRL, 6M-MFS, and 12M-CTRL respectively. Examples of positive signals are represented by white arrows. B) Glutl staining is decreased in 6M-MFS CA3 compared to 6M-CTRL. C) 12M-CTRL CA3 demonstrate decreased Glutl staining compared to 6M-CTRL, with no differences compared to 6M-MFS mice.

### Blood-brain barrier permeability is increased by age and genotype in the hippocampus

In addition, we sought to evaluate BBB permeability. Thus, Immunoglobulin (IgG) staining was utilized and evaluated in the hippocampus (**Fig. 5A, 6A, 7A**). BBB permeability, as evaluated through IgG staining, was increased in 6M-MFS (**Fig. 5B**) and 12M-CTRL mice compared to 6M-CTRL mice in the DG of the hippocampus (**Fig. 5C**), where no differences were seen between 6M-MFS and 12M-CTRL mice. Similarly, in the CA1 of the hippocampus, IgG staining was increased in 6M-MFS mice (**Fig. 6B**) and 12M-CTRL mice compared to 6M-CTRL mice (**Fig. 6C**), where no differences were seen between 6M-MFS and 12M-CTRL mice. Furthermore, in the CA3 of the hippocampus, IgG staining was increased in 6M-MFS mice (**Fig. 7B**) and 12M-CTRL mice compared to 6M-CTRL mice (**Fig. 7C**), where no differences were seen between 6M-MFS and 12M-CTRL mice.

**Figure 5.**
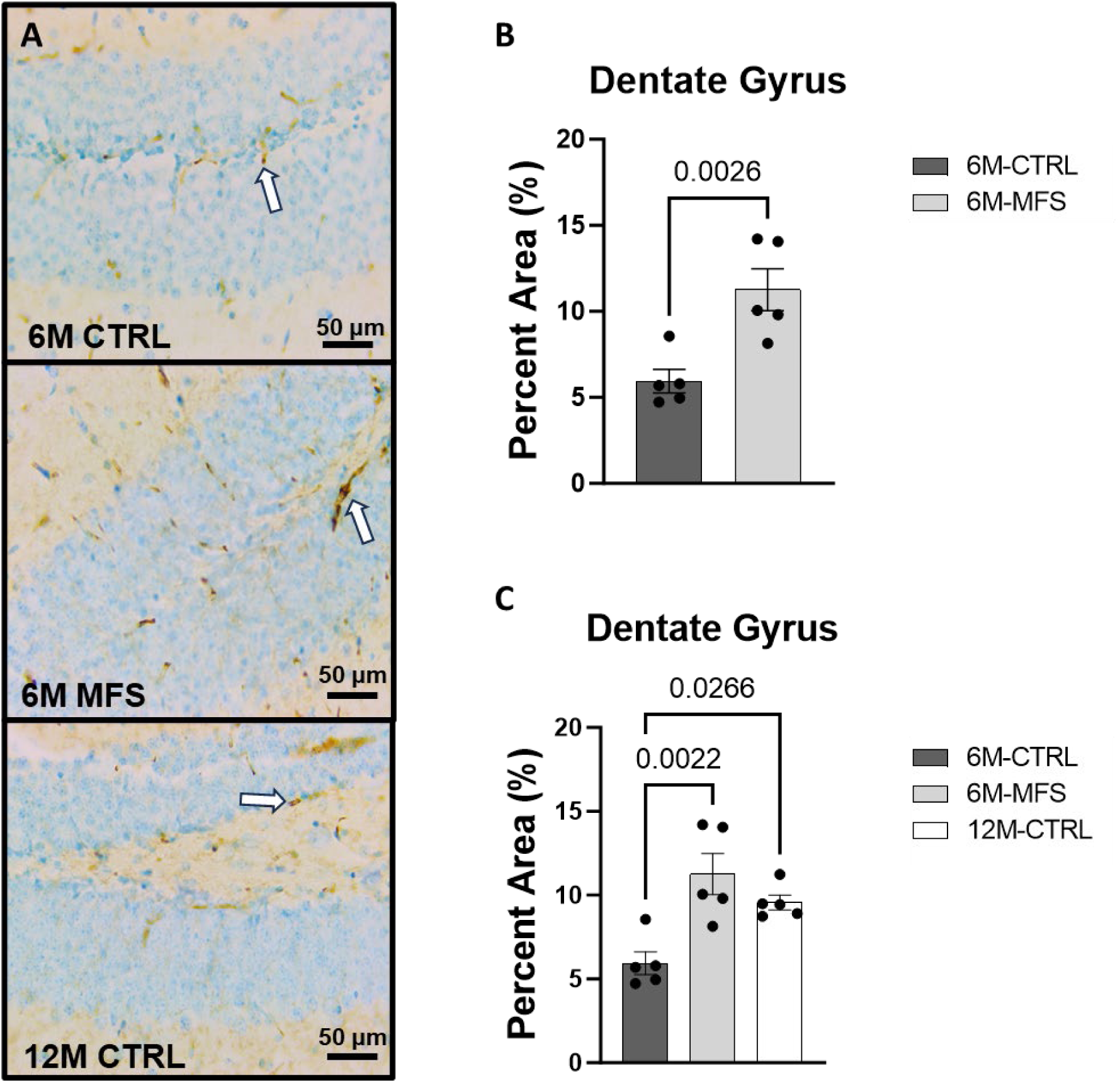
BBB permeability in the dentate gyrus (DG) of the hippocampus is increased as a function of genotype and age. A) Representative images of IgG staining in the DG of the hippocampus in 6M-CTRL, 6M-MFS, and 12M-CTRL respectively. Examples of positive signals are represented by white arrows. B) IgG staining is increased as a function of genotype, where 6M-MFS is increased compared to 6M-CTRL in the DG. C) IgG staining is increased in 6M-MFS and 12M-CTRL compared to 6M-CTRLmice in the DG. No differences were seen between 6M-MFS and 12M-CTRL.

**Figure 6.**
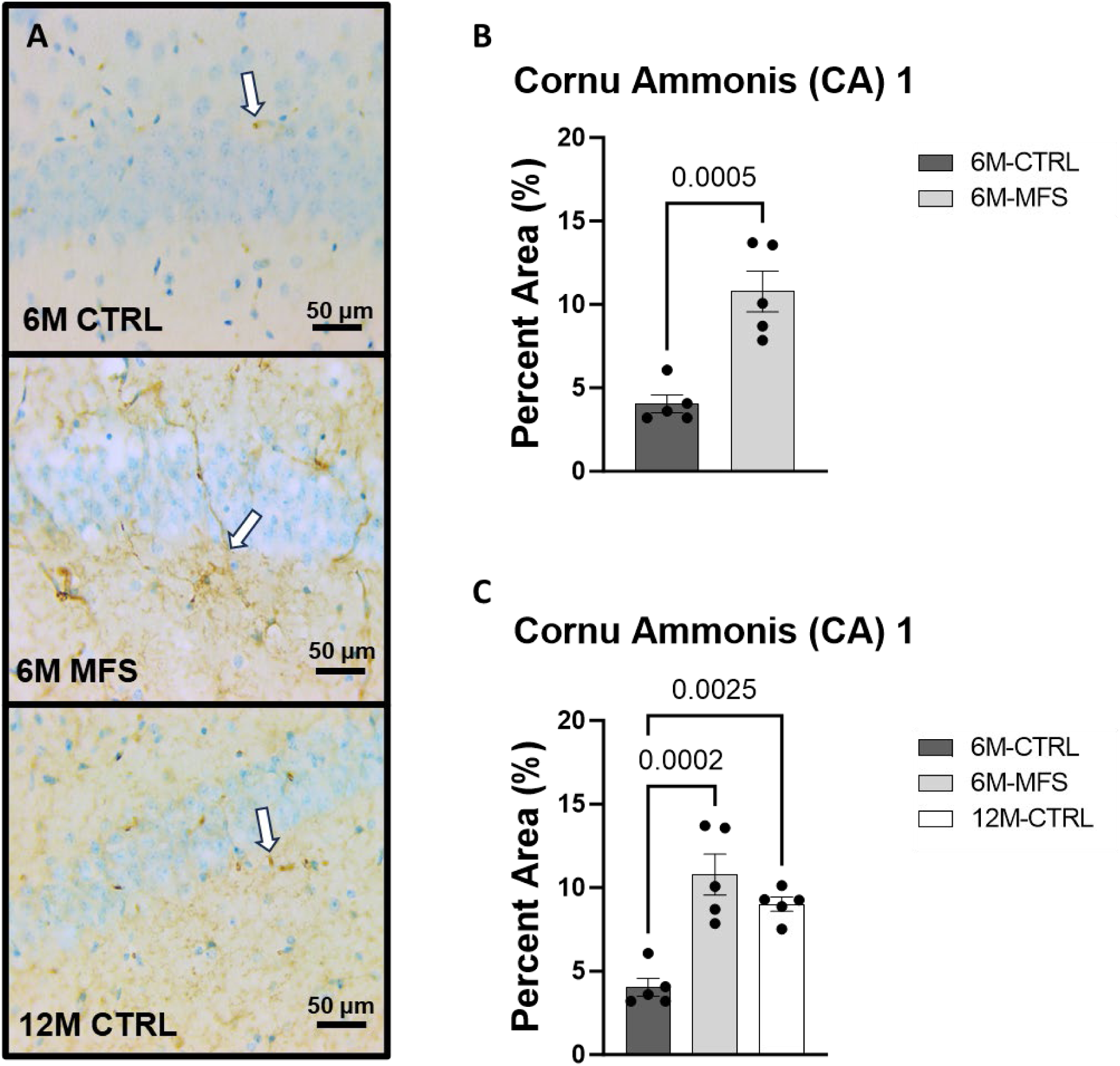
BBB permeability in the cornu ammonis 1 (CAI)of the hippocampus is increased as a function of genotype and age. A) Representative images of IgG staining in the CA1 of the hippocampus in 6M-CTRL, 6M-MFS, and 12M-CTRL respectively. Examples of positive signals are represented by white arrows. B) IgG staining is increased as a function of genotype, where 6M-MFS is increased compared to 6M-CTRL in the CA1. C) IgG staining is increased in 6M-MFS and 12M-CTRL compared to 6M-CTRL mice in the CA1. No differences were seen between 6M-MFS and 12M-CTRL.

**Figure 7.**
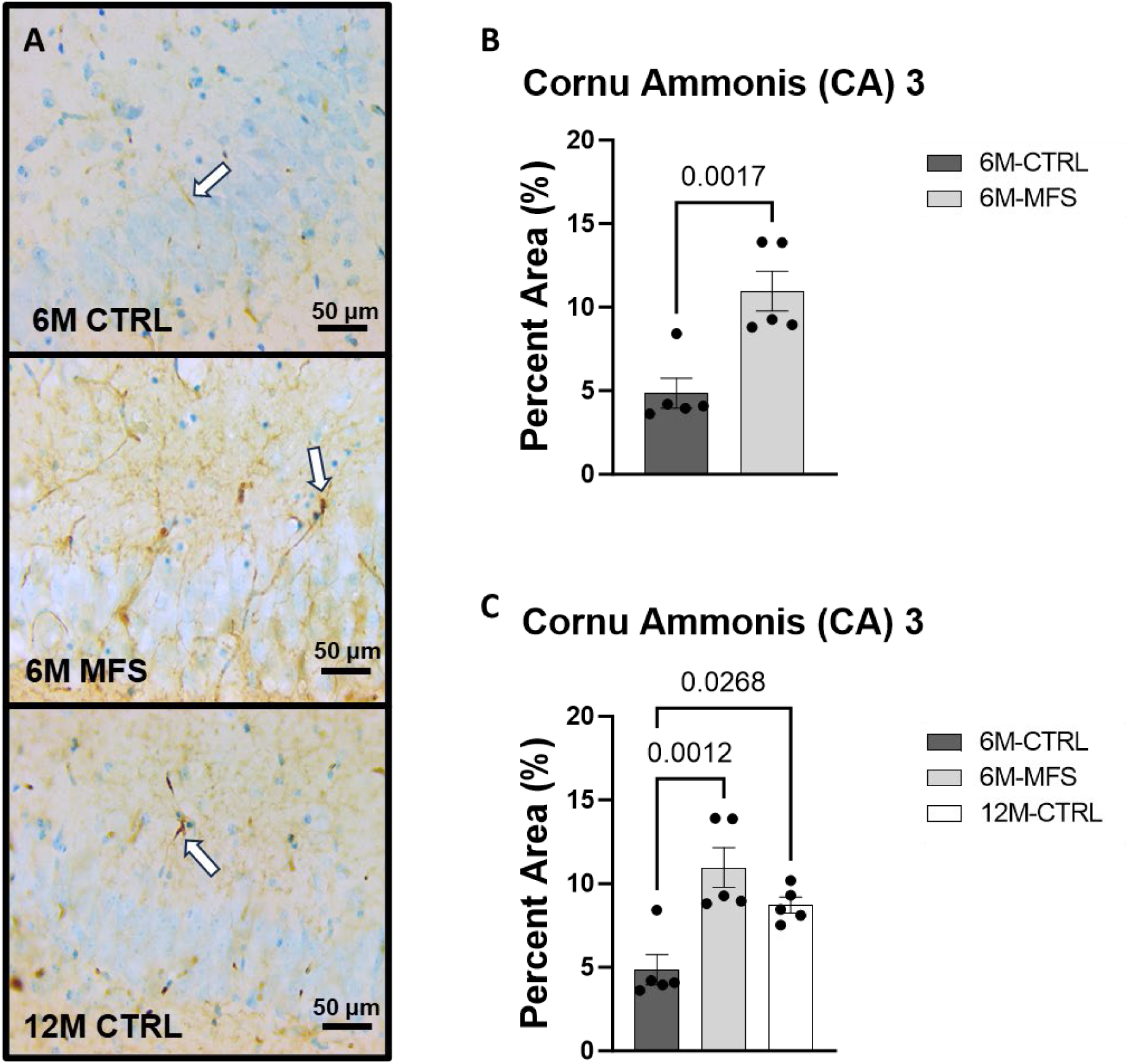
BBB permeability in the cornu ammonis 3 (CA3) of the hippocampus is increased as a function of genotype and age. A) Representative images of IgG staining in the CA3 of the hippocampus in 6M-CTRL, 6M-MFS, and 12M-CTRL respectively. Examples of positive signals are represented by white arrows. B) IgG staining is increased as a function of genotype, where 6M-MFS is increased compared to 6M-CTRL in the CA3. C) IgG staining is increased in 6M-MFS and 12M-CTRL compared to 6M-CTRL mice in the CA3. No differences were seen between 6M-MFS and 12M-CTRL.

### Microglial morphology demonstrates neuroinflammation due to age and genotype in the hippocampus

To evaluate neuroinflammation, Iba-1 staining, which stains for microglia and infiltrating macrophages, was utilized to visualize microglia soma and morphology (**Fig. 8A, 9A, 10A**). The number of microglia soma in the hippocampus (DG, CA1, and CA3), as well as the morphology of those microglia, were assessed. Previous reports by our laboratory and others have demonstrated that reactive microglia display decreased branch lengths and end points.^42,43^ Skeleton analysis was performed to quantify some of these changes in morphological features in this data set. In the DG of the hippocampus, the number of microglia soma was increased (**Fig. 8B**), while the branch length (**Fig. 8C**) and endpoint (**Fig. 8D**) per soma was decreased when comparing 6M-MFS and 6M-CTRL mice. 12M-CTRL mice were not different than 6M-MFS mice and demonstrated increased number of microglia soma (**Fig. 8E**), no difference in branch length per soma, (**Fig. 8F**) and decreased endpoint per soma compared to 6M-CTRL mice in the DG (**Fig. 8G**). In the CA1 of the hippocampus, 6M-MFS demonstrated an increase in the number of microglia soma (**Fig. 9B**), as well as a decrease in the branch lengths (**Fig. 9C**) and endpoints (**Fig. 9D**) per soma, compared to 6M-CTRL mice. Supporting the premature aging phenotype, 12M-CTRL mice demonstrated a similar increase in the number of microglia soma (**Fig. 9E**), yet without a difference in branch length per soma (**Fig. 9F**), as well as decreased endpoints per soma in the CA1, compared to 6M-CTRL mice (**Fig. 9G**). In the CA3 of the hippocampus, 6M-MFS demonstrated increased microglia soma count (**Fig. 10B**), as well as decreased branch length (**Fig. 10C**) and endpoint per microglia (**Fig. 10D**), compared to 6M-CTRL mice. Furthermore, 12M-CTRL similarly demonstrated increased microglial soma count (**Fig. 10E**), decreased branch length (**Fig. 10F**) and endpoints per soma (**Fig. 10G**), compared to 6M-CTRL mice. No differences in iba-1 staining were seen between 6M-MFS and 12M-CTRL throughout the hippocampus.

**Figure 8.**
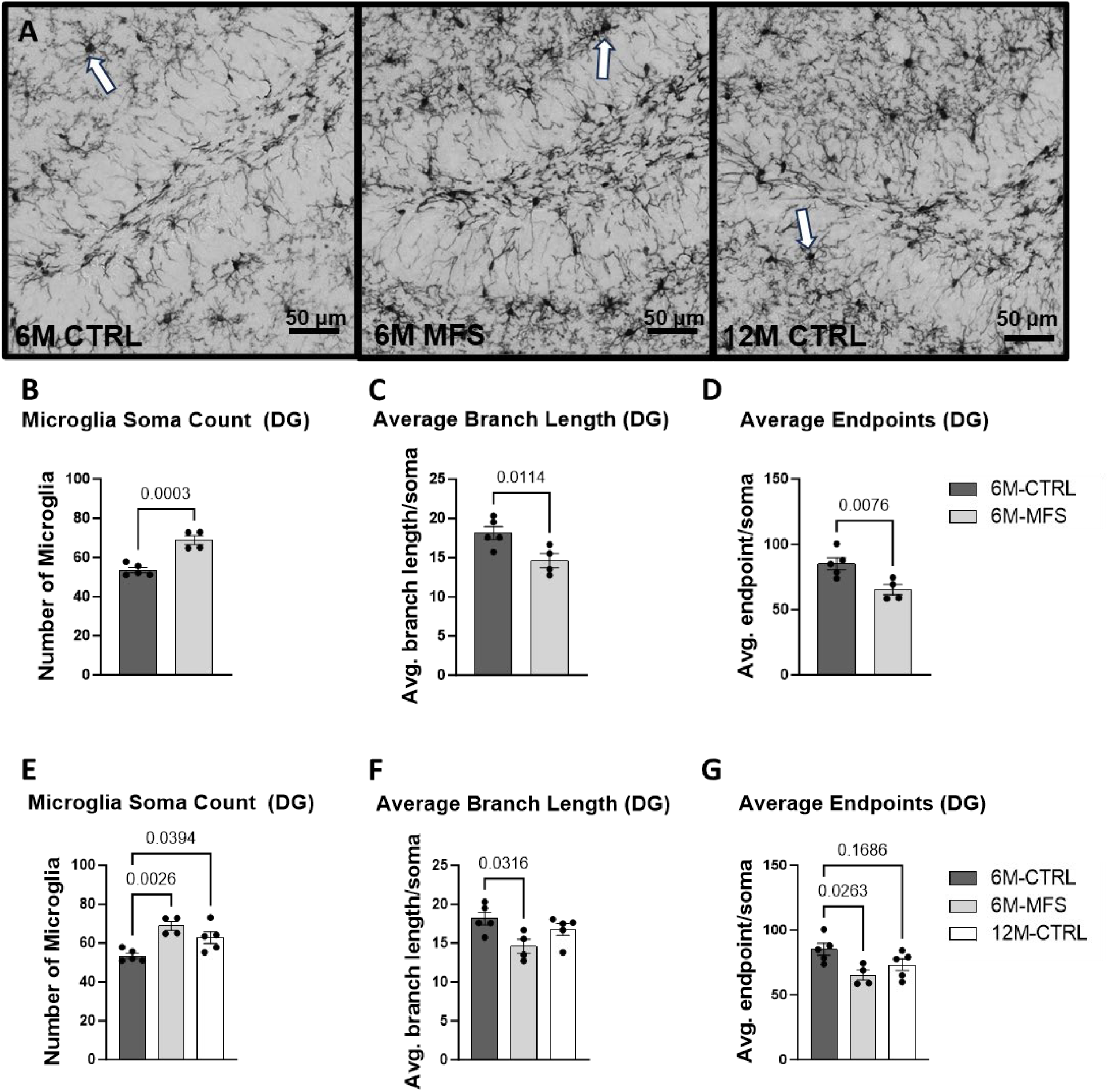
Neuroinflammation is evidenced by increased lba-1-stained soma count and decreased branch length as a function of genotype and age in the DG of the hippocampus. A) Representative images of lba-1 staining in the DG of the hippocampus in 6M-CTRL, 6M-MFS, and 12M-CTRL respectively. Examples of positive signals are represented by white arrows. 6M-MFS mice demonstrate increased B) number of microglia, C) decreased branch length, and D) decreased end point per microglia soma in the DG of the hippocampus, compared to 6M-CTRL mice. 6M-MFS and 12M-CTRLmice demonstrate increased E) number of microglia, F) 6M-MFS demonstrate decreased branch length, and G)6M-MFS and 12M-CTRLmice have decreased end point per microglia soma in the DG of the hippocampus, compared to 6M-CTRL mice.

**Figure 9.**
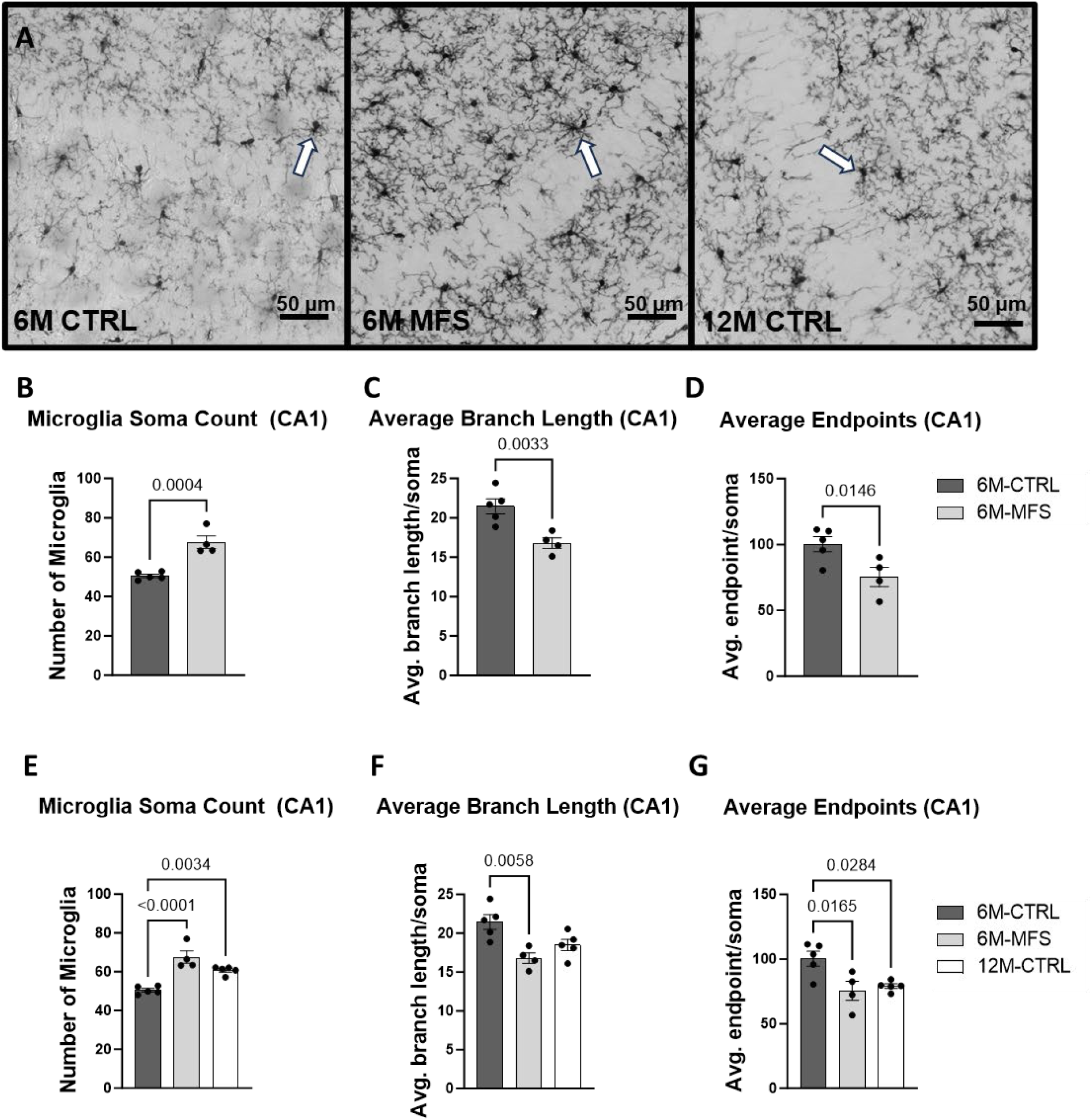
Neuroinflammation is evidenced by increased lba-1-stained soma count and decreased branch length as a function of genotype and age in the CA1 of the hippocampus. A) Representative images of lba-1 staining in the CA1 of the hippocampus in 6M-CTRL, 6M-MFS, and 12M-CTRL respectively. Examples of positive signals are represented by white arrows. 6M-MFS mice demonstrate increased B) number of microglia, C) decreased branch length, and D) decreased end point per microglia soma in the CA1 of the hippocampus, compared to 6M-CTRLmice. 6M-MFS and 12M-CTRLmice demonstrate increased E) number of microglia, F) 6M-MFS show decreased branch length, and G)6M-MFS and 12M-CTRLhave decreased end point per microglia soma in the CA1 of the hippocampus, compared to 6M-CTRL mice.

**Figure 10.**
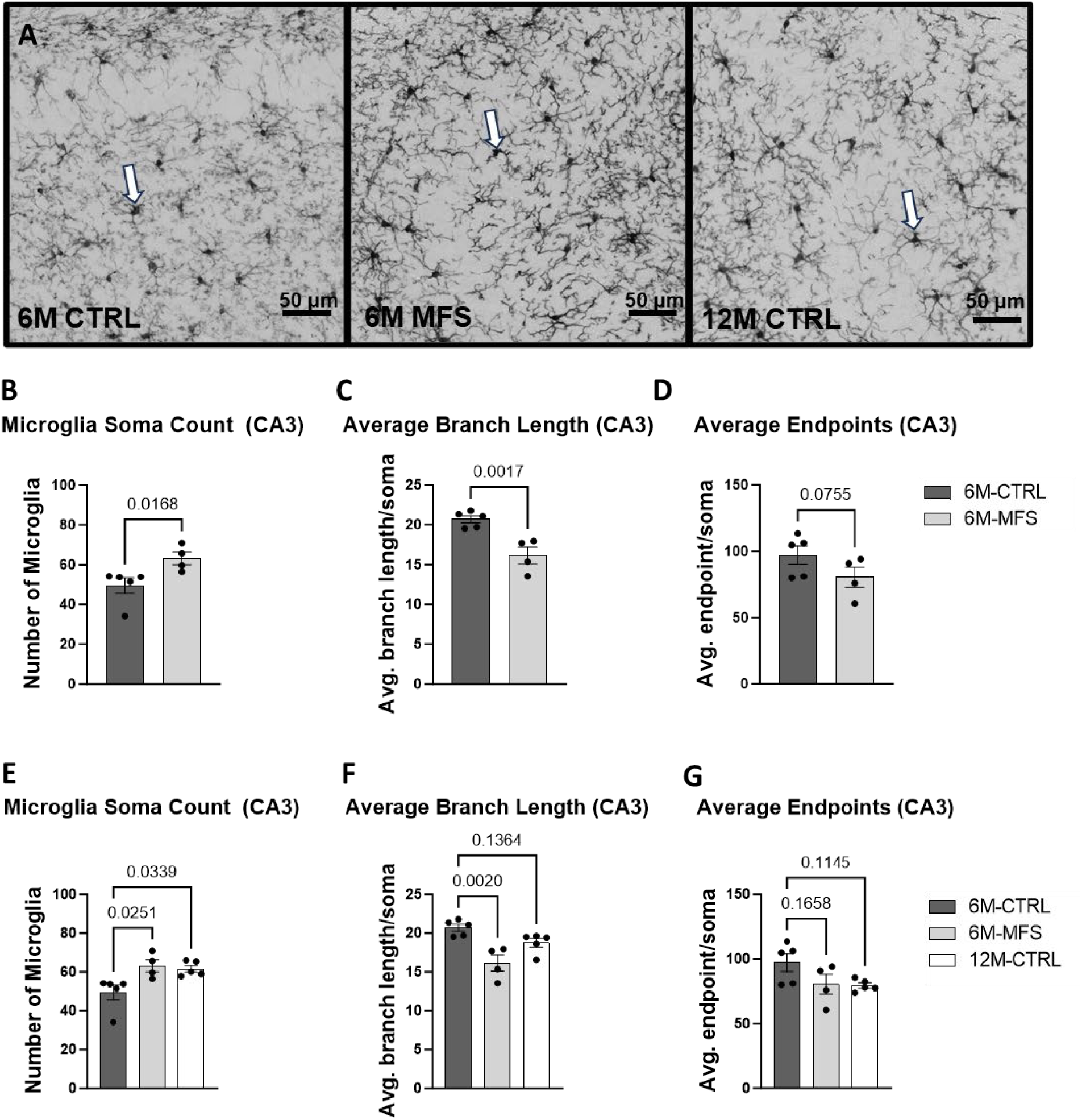
Neuroinflammation is evidenced by increased lba-1 -stained soma count and decreased branch length as a function of genotype and age in the CA3 of the hippocampus. A) Representative images of lba-1 staining in the CA3 of the hippocampus in 6M-CTRL, 6M-MFS, and 12M-CTRL respectively. Examples of positive signals are represented by white arrows. 6M-MFS mice demonstrate increased B) number of microglia, C) decreased branch length, and D) decreased end point per microglia soma in the CA3 of the hippocampus, compared to 6M-CTRLmice. 6M-MFS and 12M-CTRLmice demonstrate increased E) number of microglia, F) decreased branch length, G) decreased end point per microglia soma in the CA3 of the hippocampus, compared to 6M-CTRL mice.

## Discussion

This report investigated a well-established MFS mouse model with a demonstrated pre-mature cerebrovascular aging phenotype to evaluate corresponding neuropathology. Our findings have revealed that the *Fbn1* mutation and aging similarly contribute to decreased hippocampal microvascular density, increased BBB permeability, and neuroinflammation, supporting an MFS-associated premature aging phenotype.

Our lab investigated hippocampal microvascular density using Glut1 staining in 6M-CTRL, 6M-MFS, and 12M-CTRL mice. The data revealed a decrease in microvascular density due to the *Fbn1* mutation as well as normal aging in the DG, CA1, and CA3 areas of the hippocampus. A decrease in cerebral microvascular density, also known as cerebral microvascular rarefaction, has been shown to occur in normal aging, be accelerated in microvascular brain disorders, and play a role in reduction of perfusion.^44,45^ This reduced perfusion can further promote neurological dysfunction and tissue damage.^45^ This observed decrease in MFS mice may be correlated with TGF-β’s role in neurovascular development. TGF-β plays a crucial role in the development and differentiation of various cell types within the neurovascular unit.^46^ Utilization of a TGF-β inhibitor on human pluripotent stem cells promoted differentiation into endothelial-like cells, where observed blood-vessel-like structures began to form and promoted the synthesis of tight junctions.^46^ On the other hand, TGF-β receptor 2 (TβRII) has been demonstrated as essential for proper mediation of cerebral vascular development. Studies involving TβRII knockout mice showed an abnormal cerebral blood vessel phenotype characterized by intracerebral hemorrhage. However, this did not impact the vasculature of other major organs, indicating that TGF-β signaling is crucial for cerebral angiogenesis and development.^47^

Here, we also demonstrated increased BBB permeability in the hippocampus utilizing IgG staining methods due to both *Fbn1* mutation and age. These results are in agreement with previously reported increases in Evans Blue extravasation in the apolipoprotein E (*ApoE*)-deficient mouse model with an *Fbn1* mutation (*ApoE*^−/−^*Fbn1*^C1041G/+^).^48^ In the same study, investigators also determined that the *Fbn1* mutation was associated with decreased expression of tight junction proteins claudin-5 and occludin, as well as increased inflammatory cytokines tumor necrosis factor-alpha (TNF-α), MMPs -2/-9, and TGF-β in the choroid plexus.^48^ Therefore, it is plausible that the increased BBB permeability observed in our study can also be attributed to elevated levels of TGF-β and MMPs and disruptions in tight junctions and structural and functional integrity of BBB. The important function of TGF-β in controlling BBB function is well-established. TGF-β is one of several signaling molecules that cells of the neurovascular unit use to maintain a healthy BBB. Cells of the neurovascular unit both express and secrete TGF-β into the ECM in its latent form, where activation may be initiated by several intracellular events, including but not limited to increased reactive oxygen species (ROS), integrin activation, or protease activity.^49,50^ It has also been documented that TGF-β originating from the periphery can infiltrate the brain and activate its receptors, leading to BBB permeability.^51^ For instance, increased TGF-β signal transduction showed downregulation and destruction of tight junctions through the downstream production of MMP-2, ultimately altering the integrity of the BBB.^51^ While under normal physiological conditions, the activation of MMPs is considered a homeostatic endogenous mechanism in response to damage and injury in the BBB, under chronic conditions, this mechanism can malfunction and contribute to or exacerbate existing damage.^52^

Increased BBB permeability facilitates the extravasation of ions, proteins, glucose, glutamate, cytokines, and pro-inflammatory molecules into the extracellular space, which can promote neuroinflammation.^30,53^ In this report, both 6M-MFS and 12M-CTRL mice demonstrated increased Iba-1 positive soma counts in all three regions of the hippocampus and microglial morphology representing reactive microglia. The increased counts were accompanied by observations of cell soma hypertrophy and loss of regional heterogeneity, together, indicative of a neuroinflammatory response. As described above, *ApoE*^−/−^*Fbn1*^C1041G/+^ demonstrated increased TGF-β and MMPs in the choroid plexus, a network of blood vessels in the ventricular space of the brain.^48^ Furthermore, it has been shown that circulating TGF-β can promote neuroinflammation, where circulating TGF-β originating from the liver during liver failure was found to interact with the brain and increase neuroinflammation and neurological decline.^54^ Recent studies have confirmed that TGF-β signaling has a biphasic nature, in a manner that in response to an inflammatory stimulus in young mice, TGF-β promotes nitric oxide (NO) secretion in microglia, thus promoting a protective effect; but in aging mice, TGF-β promotes ROS secretion in microglia, thus potentiating neuroinflammation.^55^ These studies further demonstrated that the shift from NO induction to ROS induction is due to a change in the TGF-β/SMAD3 pathway, such that over availability of TGF-β actually inhibits the SMAD3 pathway.^55,56^ Therefore, it is plausible that in the mouse model of *Fbn1* mutation, chronic increased TGF-β bioavailability can contribute to the progression of pre-mature aging phenotypes and microglial activation in the brain, specifically in the hippocampus which was evaluated across three distinct regions.

To our knowledge, this study represents the first report on neuropathology assessment in the mouse model of MFS, potentially paving the way for future analyses in this domain. Throughout this report, we have identified differences in all three regions of the hippocampus, suggesting that future behavioral assays focusing on the evaluation of working memory, spatial navigation, and memory retention are warranted. This direction could provide critical insights into the cognitive impacts of MFS and guide the development of targeted therapeutic strategies.

## Study Limitations

The use of the experimental transgenic mouse model provides a systemic approach to manipulate *Fbn1* function and the downstream TGF-β signaling pathway. Changes demonstrated could be due to compensatory mechanisms of chronically upregulated TGF-β. Furthermore, due to multiple isoforms of TGF-β, investigation of these isoforms requires further evaluation. Future studies utilizing a tissue-specific approach to manipulate TGF-β signaling will allow for a better understanding of the role of TGF-β in specific cell types relevant to MFS-associated pre-mature aging of cerebral vasculature and neuropathology.

Furthermore, this study is limited to evaluating a single *Fbn1* mutation associated with MFS, specifically the mutation observed in patients who present with aortic root aneurysm. Consequently, our findings can only be directly applied to this prevalent mutation, rather than being generalizable to all individuals with MFS. This limitation underscores the need for caution when extrapolating our results to the broader MFS patient population.

It is important to emphasize that although this study incorporated both male and female subjects, it was not designed with the statistical power necessary to assess sex differences across each measured variable. Moreover, since the investigation was not aimed at exploring sex differences, factors that could contribute to variations based on sex as a biological variable, such as hormone levels, were not examined. This approach indicates that while the study acknowledges the inclusion of both sexes, it does not provide insights into how sex-specific biological factors might influence the outcomes or mechanisms under study.

## Conclusion

This study aimed to investigate the unexplored domain of neuropathological changes in connective tissue disorders, specifically MFS. In addition, comparisons were made to middle-aged CTRL mice to elucidate the effects of the premature aging phenotype, characteristic of this connective tissue disorder, on microvascular density, BBB permeability, and neuroinflammation (**Figure 11**). The findings from this research could lay the groundwork for formulating hypotheses regarding the mechanisms underlying neuropathological alterations observed in a spectrum of disorders that share a similar pathogenesis. This includes conditions characterized by chronically elevated cytokines and MMPs, such as stroke, attention deficit hyperactivity disorder (ADHD), epilepsy, repeated injuries, hypertension, and others, thereby broadening the potential impact of this study beyond MFS to a wider array of neurological conditions.

**Figure 11.**
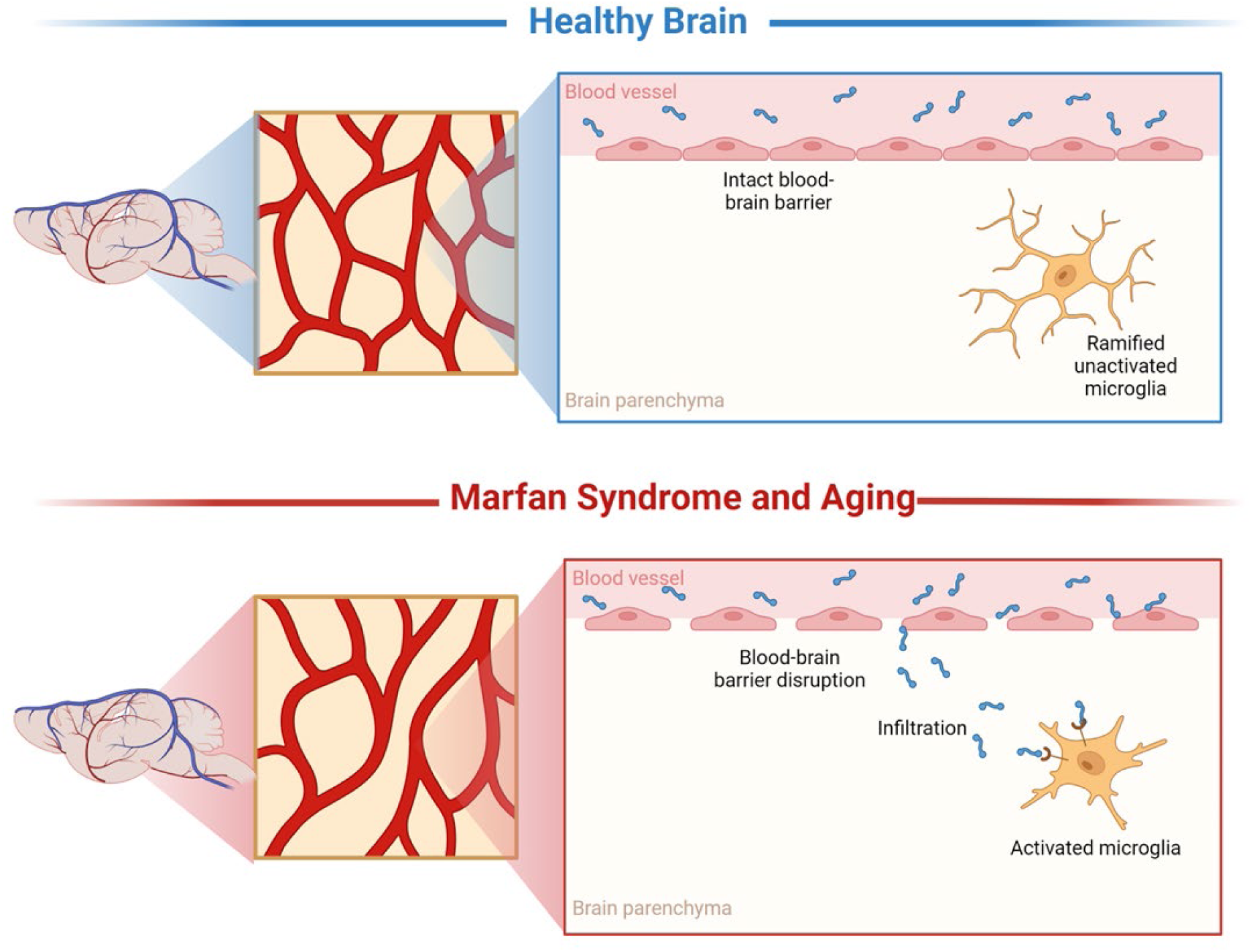
Representation of microvascular density, blood-brain barrier permeability, and neuroinflammatory alterations. These results demonstrate that MFS, similarly, to aging, supports neuropathology through decreased microvascular density, increased BBB permeability, and neuroinflammation compared to healthy controls.

## Author contributions

TMC designed experiments, collected data, analyzed data, composed figures, wrote the manuscript, and acquired funding. LPC contributed through data collection, independent analysis, and writing and review of the manuscript. ME and TCT supervised and funded the project, helped with experimental designs, reviewed the data, revised, and edited the written manuscript. All authors approved the final version of the manuscript.

## Data Availability Statement

The data sets generated for this study are available upon request to the corresponding author.

## Competing Interests

The author(s) declare no competing interests.

## Ethics Statement

Animal care was conducted according to the National Research Council Guide for the Care and Use of Laboratory Animals and the Guidelines for Animal Experiments and the Institutional Animal Care and Use Committee. All experiments were in compliance with ARRIVE guidelines. The animal study was reviewed and approved by the Institutional Animal Care and Use Committee at Midwestern University-Glendale, Arizona [IACUC protocols AZ-3006, AZ-2989, & AZ-2936].

## Funding

This project was funded, in part, by a National Institutes of Health (R36AG083385) to T.M.C, Valley Research Partnership P1a grant (2232011) to T.M.C, National Institutes of Health (R15HL145646) to M.E., National Institutes of Health (R01NS100793) to T.C.T, and Phoenix Children’s Hospital Mission Support. The content is solely the responsibility of the authors and does not necessarily represent the official views of the National Institutes of Health, Valley Research Partnership, or Phoenix Children’s Hospital.

## Notes

### Competing Interest Statement

The authors have declared no competing interest.

